# The *Onychomys* pangenome reveals the unique molecular adaptations that confer toxin resistance

**DOI:** 10.1101/2025.10.28.685014

**Authors:** Claudia Perez Callez, Jingtao Lilue, Elizabeth Anderson, Leanne Haggerty, Fergal J. Martin, Shane A. McCarthy, David J. Adams, David Thybert, Ashlee H. Rowe, Thomas M. Keane

## Abstract

Novel traits enable many rodents to thrive in extreme environmental niches. Predatory grasshopper mice (*Onychomys* sp.) have co-evolved resistance to painful and lethal neurotoxins produced by their scorpion prey. Previous work reported that grasshopper mice have structural and functional modifications in sodium channel Nav1.8 that block the effect of painful toxins. However, key questions remain about the molecular adaptations underlying toxin resistance. We produced the first high-quality reference genomes and annotations for *Onychomys* species and *Peromyscus eremicus*. We implemented a comprehensive pipeline to detect positive selection across genome-scale datasets and identified *Onychomys-specific* mutations in Nav1.3 (*Scn3a),* a sodium channel gene expressed in the central nervous system and the peripheral sensory system after nerve damage. We detected an *Onychomys*-specific tandem gene duplication of the *Cblif* gene, which encodes a glycoprotein crucial for vitamin B12 absorption. This adaptation likely supports the species’ dietary specialisation and modified stomach morphology, where parietal cells expressing *Cblif* are especially numerous. Our study provides a key step for establishing the *Onychomys* species as a model system for studying toxin resistance, alternative pain phenotypes, and behavioural traits related to predator-prey interactions.

## Introduction

The order Rodentia underwent an extraordinary adaptive radiation during the Cenozoic era and accounts for nearly half of all known mammalian diversity, containing over 2,000 species^1,2^. There are several examples of how rodents have adapted and thrived in environmental niches, resulting in novel traits not found in laboratory mice^3,4^. One example is the *Onychomys* species (grasshopper mice) (Figure 1A), which have co-evolved resistance to several venomous toxins that cause injury to other mammals and humans^5^. There are three grasshopper mice species: *Onychomys torridus* (southern), *Onychomys leucogaster* (northern), and *Onychomys arenicola* (Mearns or Chihuahuan), found in short-grass prairies, shrub deserts, and desert grasslands of the western United States and Northern Mexico (Figure 1B and C). Hybridisation is extremely rare in *Onychomys;* however, there is evidence for occasional hybridisation between *O. arenicola* and *O. leucogaster*^6^. Grasshopper mice are nocturnal rodents and obligate carnivores, with arthropods making up most of their diet^7^. Consequently, they present differences in behaviour, morphology, and physiology. For example, grasshopper mice are highly aggressive, attacking almost any moving object that is not significantly larger than themselves; their jaws have evolved to provide more forceful bites; their stomachs are adapted to deal with exoskeletons, and their claws are modified to enhance prey capture^8^.

**Figure 1:**
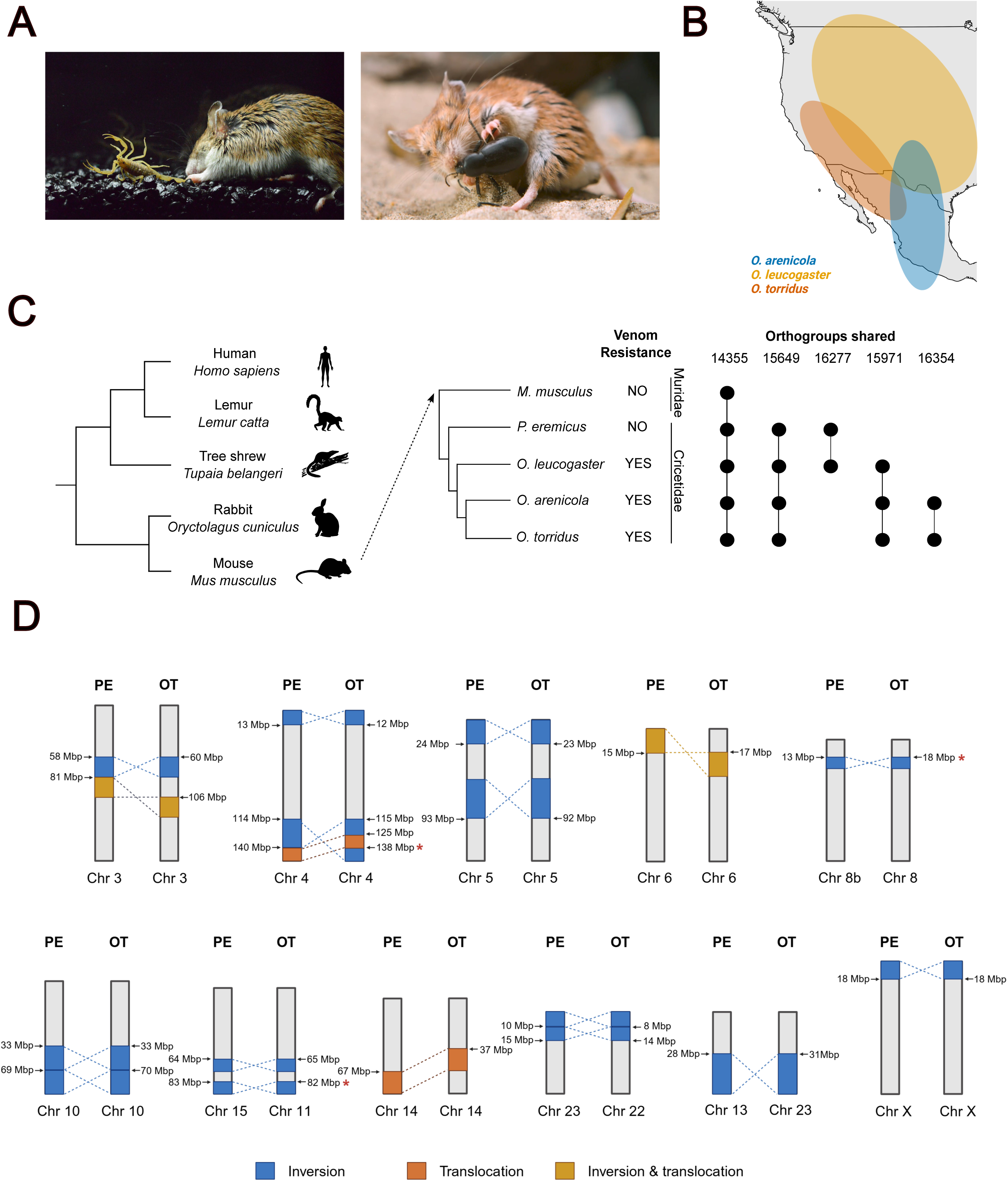
The *Onychomys* species and pangenome. **A)** Left: Grasshopper mouse attacking an Arizona bark scorpion (*Centruroides sculpturatus*). Image courtesy of A. Rowe and M. Rowe. Right: Grasshopper mouse attacking a pinacate beetle (*Eleodes*). Image courtesy of Ann Prum, Cone Flower Studios, and A. Rowe and M. Rowe. **B)** Geographic distribution of *Onychomys* species in North America: *O. arenicola* (blue), *O. leucogaster* (yellow), and *O. torridus* (orange). **C)** Euarchontoglires mammals phylogeny highlighting *Onychomys* species and their closest outgroup (*Peromyscus eremicus*), with a focus on the phenotype of interest: venom resistance. Tree is not to scale. On the right, the number of orthogroups shared within several combinations of rodents is displayed. **D)** Major synteny breaks between *Peromyscus eremicus* (PE) and *Onychomys torridus* (OT) chromosomes (**\***indicates rearrangements not supported by *O. arenicola* and *O. leucogaster*). Created with BioRender.com.

Many desert arthropods, including scorpions, centipedes, spiders, and pinacate beetles, are chemically defended. These animals use their toxic stings, bites, and sprays to subdue prey and deter predators. However, grasshopper mice have evolved resistance to the chemical weapons of their prey. For example, they are resistant to the venom of scorpions in the genus *Centruroides*, commonly known as bark scorpions in some regions of the southwest. Species in the genus *Centruroides* produce venoms containing toxins that target sodium (Na+) and potassium (K+) ion channels in nerve and muscle tissue causing pain and neuromuscular dysfunction^9–11^. Geographic patterns of grasshopper mouse resistance covary with the chemical defenses of their local *Centruroides* species, consistent with an evolutionary arms race. *O. torridus* is sympatric with *Centruroides sculpturatus* (AZ bark scorpion) in the Sonoran desert, and *O. arenicola* is sympatric with *Centruroides vittatus* (striped bark scorpion) in the Chihuahuan desert. Both scorpion species produce toxins that target Na+ and K+ channels in nerve and muscle tissue, yet *C. sculpturatus* is more toxic and painful than *C. vittatus.* While *O. torridus* and *O. arenicola* are each resistant to the toxins of their local scorpion species, they exhibit only moderate resistance to the toxins from non-local *Centruroides* species^5,12^. Arizona bark scorpion venom is not only lethal to small mammals, the neurotoxins can kill a human infant or young child^13,14^. Lethal toxins bind the skeletal muscle voltage-gated sodium (Nav) channel Nav1.4 to block muscle contraction, causing asphyxiation. However, *O. torridus* express amino acid substitutions in their muscle sodium channel Nav1.4 that reduce the effects of bark scorpion venom on muscle contraction^15^. In contrast, pain-inducing toxins target Nav channels in sensory neurons that transmit pain signals to the brain. Grasshopper mice have structural modifications in sodium channel Nav1.8, that enable the channel to bind venom toxins, which inhibit Na+ currents and block action potential propagation^16^. Essentially, toxins act as analgesics to block pain signals.

In addition to feeding on scorpions, grasshopper mice also prey on pinacate beetles, which spray toxic aerosols in the face of their predators during attacks^17^. Pinacate beetle sprays are acidic mixtures containing benzoquinones that target the sensory transduction channels TRPA1 and TRPV1 (TRP, transient receptor potential), causing irritation of the eyes and nasal epithelium^18–21^. While observations of interactions between grasshopper mice and pinacate beetles suggest that the mice have evolved resistance to pinacate beetle sprays, the mechanism of resistance is unknown.

Collectively, the studies described above suggest that grasshopper mice have evolved a complex physiological phenotype comprising resistance to both painful and lethal chemical weapons targeting distinct Nav and TRP channels that regulate sensory and muscle function. However, toxin resistance across multiple physiological systems raises critical questions regarding modifications to additional receptors and signaling pathways involved in sensory and neuromuscular function. To address these questions, we developed a comparative genomics framework to understand the genetic basis for adaptive toxin resistance. While the majority of studies to date focused on *O. torridus*, there are two other *Onychomys* species (*O. leucogaster* and *O.arenicola*) that are less resistant to the bark scorpion venom^5^. The closest outgroup to *Onychomys* is *Peromyscus*^22,23^*. Peromyscus eremicus*, the cactus mouse, coexists with *O. torridus* in the Sonoran Desert. While *P. eremicus* are known to consume some insects, there are no reports of *P. eremicus* consuming scorpions or pinacate beetles. *P. eremicus* is not known to be resistant to either bark scorpion venom or pinacate beetle spray. We created reference genomes for the three *Onychomys* species and the closest outgroup, *Peromyscus eremicus*. We carried out positive selection analysis at the genome scale at two levels: within and across *Onychomys* species, and combined with population sequencing and RNA-seq data to determine fixation in the population. In parallel, we characterised the molecular response to toxin exposure by analysing differential gene expression (RNA-seq before and after pinacate beetle toxin exposure) in pain-signalling tissues.

## Results

### Chromosome-scale *Onychomys* genomes

Chromosome-scale de novo assemblies of *Onychomys arenicola, Onychomys leucogaster, Onychomys torridus,* and *Peremyscus eremicus* were generated using a combination of PacBio continuous long reads (CLR) and HiFi sequencing, and Hi-C for chromosome scaffolding (Table 1). The genomes underwent several rounds of manual curation, polishing, and base error correction (see methods), resulting in assemblies of between 2.40 (*P. eremicus* H2) to 2.69 Gbp (*O. arenicola* H1) for haplotypes with an X chromosome (excluding unknown bases, e.g. Ns). We improved an earlier *O. torridus* genome assembly produced from CLR reads by aligning against the *O. arenicola* and *O. leucogaster* genomes. We concluded that chromosomes 3, 9, 22, and X had misassemblies due to scaffolds in the incorrect order, which were subsequently corrected (Supplementary Table 1). Pan-metazoan BUSCO gene content completeness ranged from 92.80-94.80% across the assemblies (Table 1). Approximately 0.06-0.7 Gbp of sequence is unplaced per strain, and gaps (N bases) consist of 0.32-5.30 Mbp (*O. arenicala*, *O. leucogaster*, and *P. eremicus*) and 244.31 Mbp (*O. torridus*). A foreign DNA contamination check identified a *Sarcocystis neurona* (neural parasite) genome in the *O. arenicola* genome. The other genomes did not present any significant contaminants.

**Table 1:** Summary statistics of the *Onychomys* and *Peromyscus* reference genomes (chromosomes 1-23 and X; except *Onychomys leucogaster* H1, which does not contain chromosome X).

The total repeat content of these genomes ranges as follows: SINEs (213.37-222.03 Mbp), LINEs (290.28-334.35 Mbp) and ERVs (357.97-379.25 Mbp). We compared the fraction of the genome consisting of repeats with all available high-quality rodent genomes (long reads and chromosome-level). Whilst the fraction of the genome consisting of SINEs is similar in Muridae (5.13-9.23%) and Cricetidae (6.75-8.91%), LINEs and ERVs show significant differences. ERVs occupy 11.91-14.95% in the Cricetidae compared to 7.69-12.21% in Muridae (t=7.82, p < 0.0001). LINE elements occupy 10.6-17.47% in the Cricetidae compared to 12.64-22.47% in Muridae (Wilcoxon W=11, p=0.0009) (Supplementary Figure 1 and Supplementary Table 2).

Gene annotations were produced by the Ensembl Gene Annotation pipeline. The primary annotations were generated by aligning short-read transcriptomic datasets, sourced from the European Nucleotide Archive (ENA), to the respective genomes. To address regions where transcript-based evidence was incomplete, additional annotation support was incorporated by aligning mammalian SwissProt proteins from UniProt^24^. Complementary evidence was also provided by mapping mouse coding sequences from the GENCODE resource^25^ through pairwise genome alignments. The total number of protein-coding genes ranges from 19,018-20,373 across the *Onychomys* species.

### Sequence diversity in the *Onychomys* pangenome

We performed a whole-genome alignment for *O. torridus* and *P. eremicus,* revealing 11 chromosomes with structural variants exceeding 1Mbp, and a total number of 452 inversions and 24 rearrangements larger than 1 Mbp (Figure 1D, Supplementary Figure 2 and Supplementary Table 3). All the rearrangements were supported by the *O. arenicola* and *O. leucogaster* genomes, except for the inversion in chromosome 8b, the last inversion in chromosome 4, and the last inversion in chromosome 15. In contrast, we detected 674 inversions between *O. leucogaster* and *O. torridus* (8 bigger than 1 Mbp), 698 between *O. arenicola* and *O. torridus* (10 bigger than 1 Mbp) and 574 between *O. leucogaster* and *O. arenicola* (15 bigger than 1 Mbp) (Supplementary Table 4).

To detect sequence variation and conservation within the *Onychomys* species, we used minigraph^26^ to build an *Onychomys* species pangenome. In this graph, *O. torridus* serves as the backbone, with non-reference sequences represented as branches in the graph (Figure 2A). Since the pangenome graph fully represents the genomes of the species, we can use it to interrogate the complete set of *Onychomys* sequences and haplotypes. By traversing the pangenome graph, we identified all non-*O. torridus* paths and merged these into the most divergent non-reference loci (see Materials and Methods). Figure 2A shows the amount of non-reference sequence across each chromosome for the *Onychomys* species, which varies between 54.8 Kbp per 1 Mbp in chromosome 20 up to 86.7 Kbp per 1 Mbp on chromosome X (Supplementary Table 5). The non-reference regions in the pangenome vary in size between 50 bp (default minimum threshold of minigraph) and 454.8 Kbp (Figure 2A). The *Onychomys* genomic loci of highest divergence (top 5%) contain 2,262 protein-coding genes. We examined the protein classes of these genes using PANTHER, and the results show an enrichment for defence, immunity and dopamine biosynthetic protein classes (Figure 2B). One of the protein classes that was enriched was the *Clec4* gene family on chromosome 3 (Figure 2A). The *Clec4* gene family encodes proteins containing C-type lectin domains, which contribute to various immune functions such as pathogen recognition, cell adhesion, and regulation of inflammatory responses^27^.

**Figure 2:**
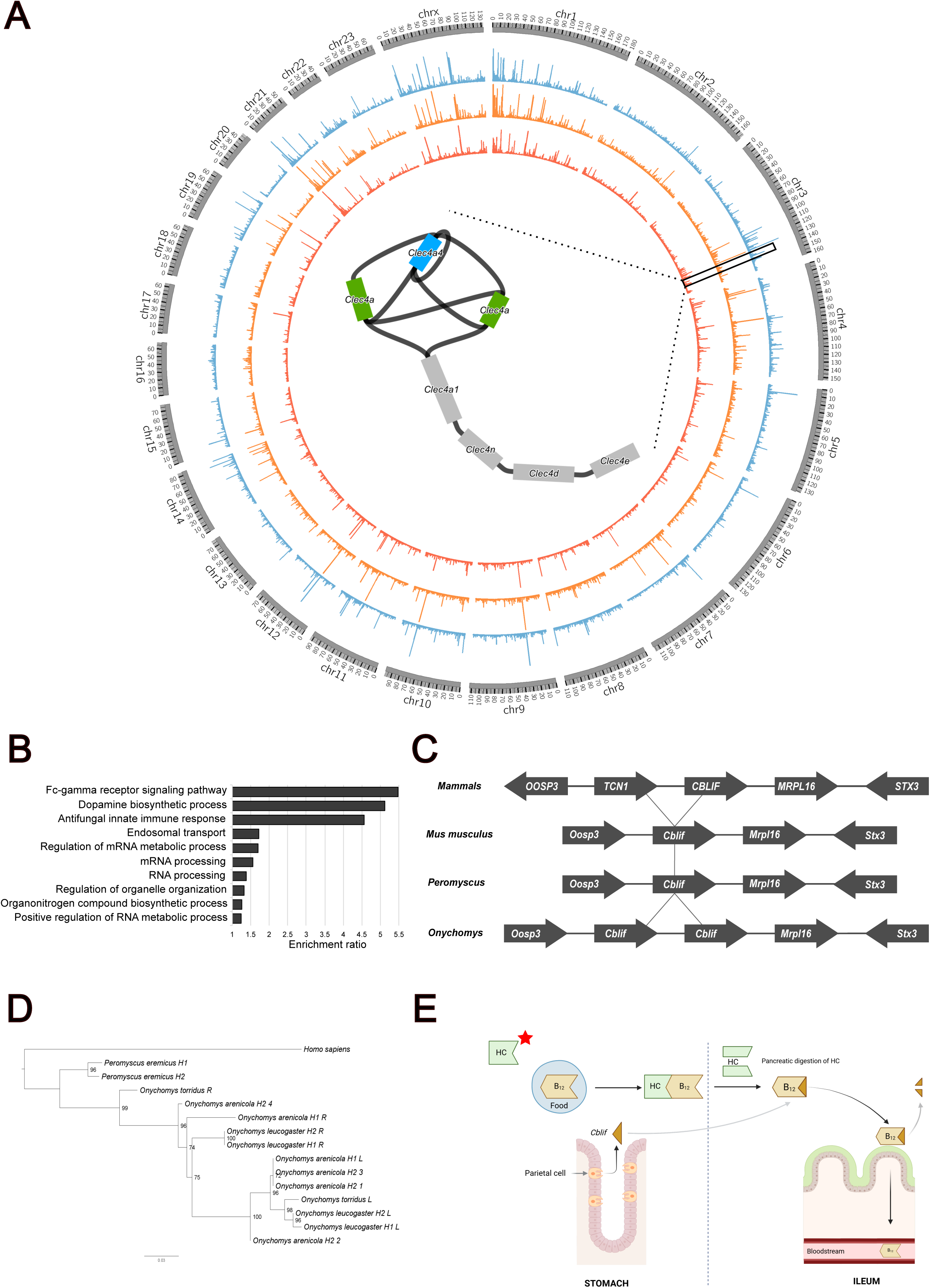
The *Onychomys* pangenome. **A)** *Onychomys torridus* chromosomes arranged with histogram bars that represent the amount of non-reference sequence in the other *Onychomys* species, ranging from 0.8-200 Kbp. *O. arenicola* (blue), *O. leucogaster* (orange), and *O. torridus* (red). The central region displays an assembly graph of the *Clec* gene family; blue indicates a duplication in *O. arenicola* and green a rearrangement. **B)** Panther overrepresentation test showing enriched gene functions among the top 5% most divergent genomic loci. **C)** Comparative gene structures of *Cblif* and *TCN1A* in *Homo sapiens, Mus musculus, Peromyscus, and Onychomys*. *Tcn1a* is absent in Muridae and Cricetidae; *Cblif* is duplicated in *Onychomys*. **D)** Phylogenetic tree of *Cblif* sequences, showing haplotypes and chromosomal location of each copy (R: right; L: left). In *O. arenicola* H2, four copies are present, with copy 1 at the far right and copy 4 at the far left. **D)** Vitamin B12 absorption pathway. HC (star) denotes haptocorrin. Created with BioRender.com.

We detected an *Onychomys*-specific tandem gene duplication of the *Cblif* gene (Figure 2C), which encodes a glycoprotein crucial for vitamin B12 absorption^28^. All the *Oychomys* species contain at least two copies of the *Cblif* gene, and *O. arenicola* appears to be in a heterozygous state with two and four copies on each haplotype. A phylogenetic tree for the *Cblif* gene, incorporating sequences from *P. eremicus* and *Homo sapiens*, clusters the copies into two monophyletic groups by chromosome order, suggesting an *Onychomys* ancestral duplication event at this locus (Figure 2D).

### Positive selection

We implemented a comprehensive pipeline to detect positive selection across genome-scale datasets to identify genes under evolutionary pressure that may contribute to the phenotypic or functional differences found in the *Onychomys* species (Figure 3A). We ran the test at two levels: within *Onychomys* species and across a set of 14 mammalian species (Supplementary Table 6). For the *Onychomys* species, we designated the ancestral branch as the foreground branch to test for positive selection. Initially, we analysed 10,769 genes, which were reduced to 314 after running PAML^29^, and further narrowed down to 74 genes following a false discovery rate (FDR) analysis. For the mammalian species analysis, we marked each *Onychomys* species as the foreground branch. When *O. torridus* was the foreground branch, we started with 11,261 genes, which decreased to 317 after multiple testing corrections (Figure 3B). For *O. arenicola* and *O. leucogaster*, we initially had 11,154 and 11,128 genes, respectively, which were subsequently narrowed down to 198 and 165 (Supplementary Table 7). Among the genes highlighted as positively selected, a significant portion were false positives due to alignment errors; consequently, the number of high-quality candidates was 11, 13, 5 and 2 for the foreground branches *Onychomys* ancestral, *O. torridus*, *O. arenicola*, and *O. leucogaster*, respectively (Table 2).

**Figure 3:**
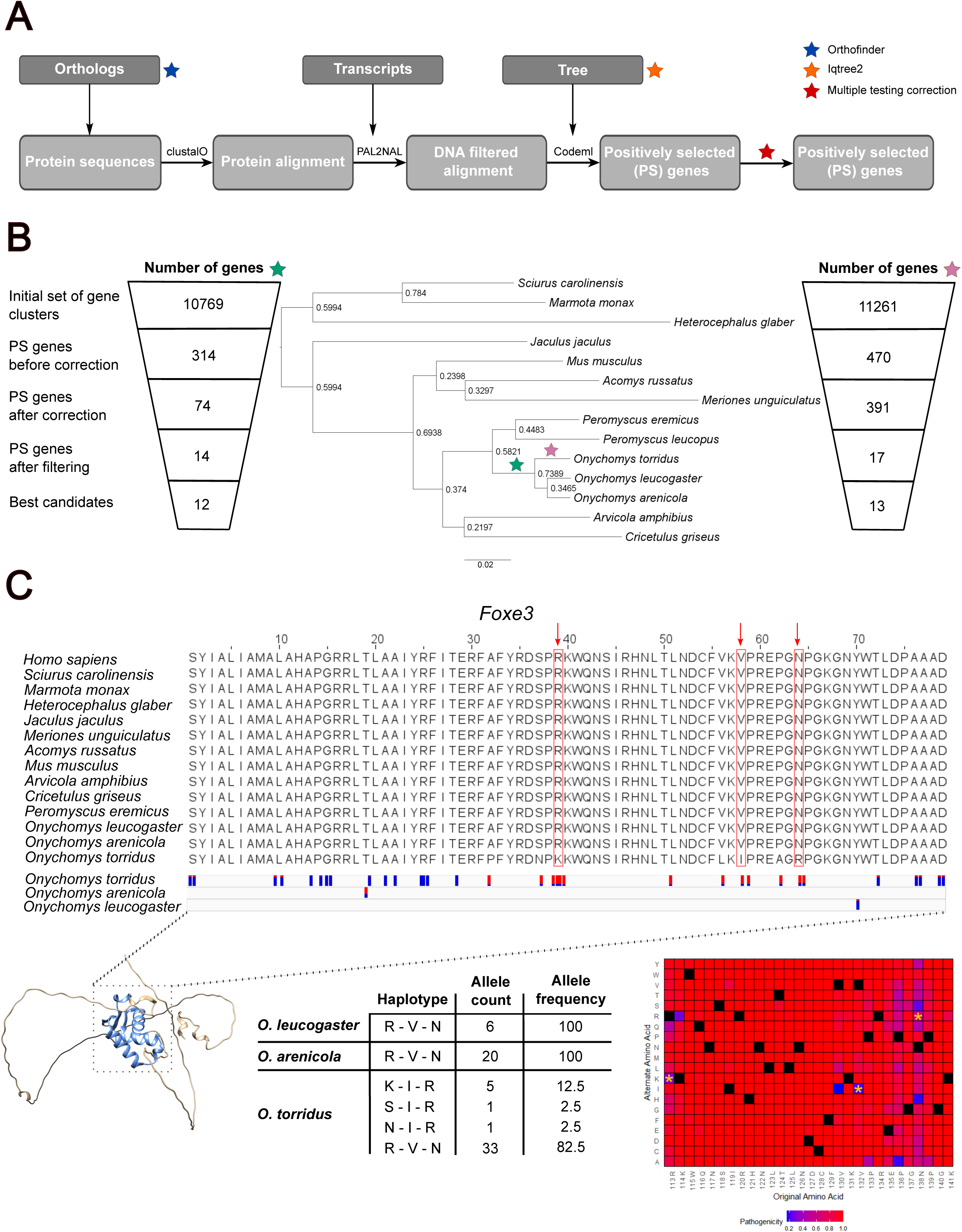
***Onychomys* positive selection analysis. A)** Schematic of the positive selection analysis pipeline. **B)** Number of genes retained at each step of the pipeline, including filtering, for the *Onychomys* ancestral branch (left; green star in the species tree) and the *Onychomys torridus* ancestral branch (right; purple star). **Centre**: species tree generated by OrthoFinder using the STAG algorithm and rooted with STRIDE, showing all species included in the analysis. **C)** Example of a positively selected gene in *O. torridus: Foxe3*. **Top**: multiple sequence alignment showing the forkhead domain; red boxes and arrows indicate sites under positive selection. Below the alignment, population data show the allele fraction histogram across Onychomys species - in red, the alternate allele and in blue the reference allele. **Middle**: table summarising haplotypes, allele counts, and allele frequencies across *Onychomys* species. **Bottom left**: AlphaFold-predicted 3D structure of *O. torridus Foxe3*, with the forkhead domain highlighted. **Bottom right**: AlphaMissense heatmap (based on the human ortholog), predicting the pathogenicity of all possible amino acid substitutions; asterisks mark substitutions found in *O. torridus*.

**Table 2:** Genes under positive selection in the *Onychomys* species. Genes with substitutions found in highly conserved regions are in bold. **indicates not significant when using a four-fold degenerate site tree before multiple testing correction. *Only significant when using a four-fold degenerate site tree after multiple testing correction.

In the analysis where *O. torridus* was the foreground branch, *Foxe3* was a notable positively selected gene. This gene encodes a forkhead transcription factor essential for lens vesicle closure and epithelial cell proliferation^30–32^. Three residues, Lys[107], Ile[126], and Arg[132], were identified as positively selected (Figure 3C). In *O. torridus*, the amino acid substitutions Lys[107] and Ile[126] were predicted by AlphaMissense^33^ to be non-pathogenic in *H. sapiens*; in contrast, Arg[132] was predicted to be pathogenic, with a score of 0.614, although only slightly above the threshold for uncertainty (0.564).

We generated short-read population sequencing data for the three *Onychomys* species. *O. torridus* samples were collected from two populations: the SRER (Santa Rita Experimental Range) Arizona; and the Chiricahua Mountains, Arizona. *O. arenicola* were collected from the Organ Mountains, New Mexico. *O. leucogaster* were collected from a sand sage prairie habitat, Kansas. *P. eremicus* were collected from the Santa Rita Experimental Range, Arizona. We integrated the candidate genes with the population sequencing data to validate whether the detected mutations were fixed or varying across the population. We generated Illumina sequencing data for 22 *O. torridus*, 10 *O. arenicola,* and 4 *O. leucogaster* individuals (Supplementary Table 8). Interestingly, in the case of *Foxe3*, all individuals in the *O. torridus* population were either homozygous for the ancestral allele or heterozygous (Figure 3C). All of the *O. arenicola* and *O. leucogaster* individuals are homozygous for the ancestral allele, and the mutations are significantly enriched in the *O. torridus* individuals (Fisher’s exact test, two-sided p-value = 0.027). Genomewide heterozygosity rates are similar across the *Onychomys* species (*O. torridus* = 0.623%, *O. arenicola* = 0.371%, and *O. leucogaster* = 0.779%), suggesting that the heterozygote allele observed in *O. torridus Foxe3* may be functional.

### Sodium channel - *Scn3a*

It is known that structural and functional modifications in the sodium channel Nav1.8 decrease the effect of toxins^16^. When testing for positive selection in the ancestral *Onychomys* branch, we identified positive selection in *Scn3a* (Nav1.3), which encodes a sodium channel alpha subunit (Figure 4C, D). Two residues, Ser[867] and Ser[868], were identified as being under positive selection, and the second one is predicted to be phosphorylated (Figure 4A). Notably, these amino acid changes in *Onychomys* were predicted by AlphaMissense to be non-pathogenic in *Homo sapiens* (Figure 4B). *Scn3a* is a transmembrane protein, and the structure and location of the residues positively selected are found in the S5-S6 loop of DII, shown in Figure 4D. For *Scn3a*, no variants were detected in the population data, suggesting that these changes may be fixed in all species in the *Onychomys* lineage.

**Figure 4:**
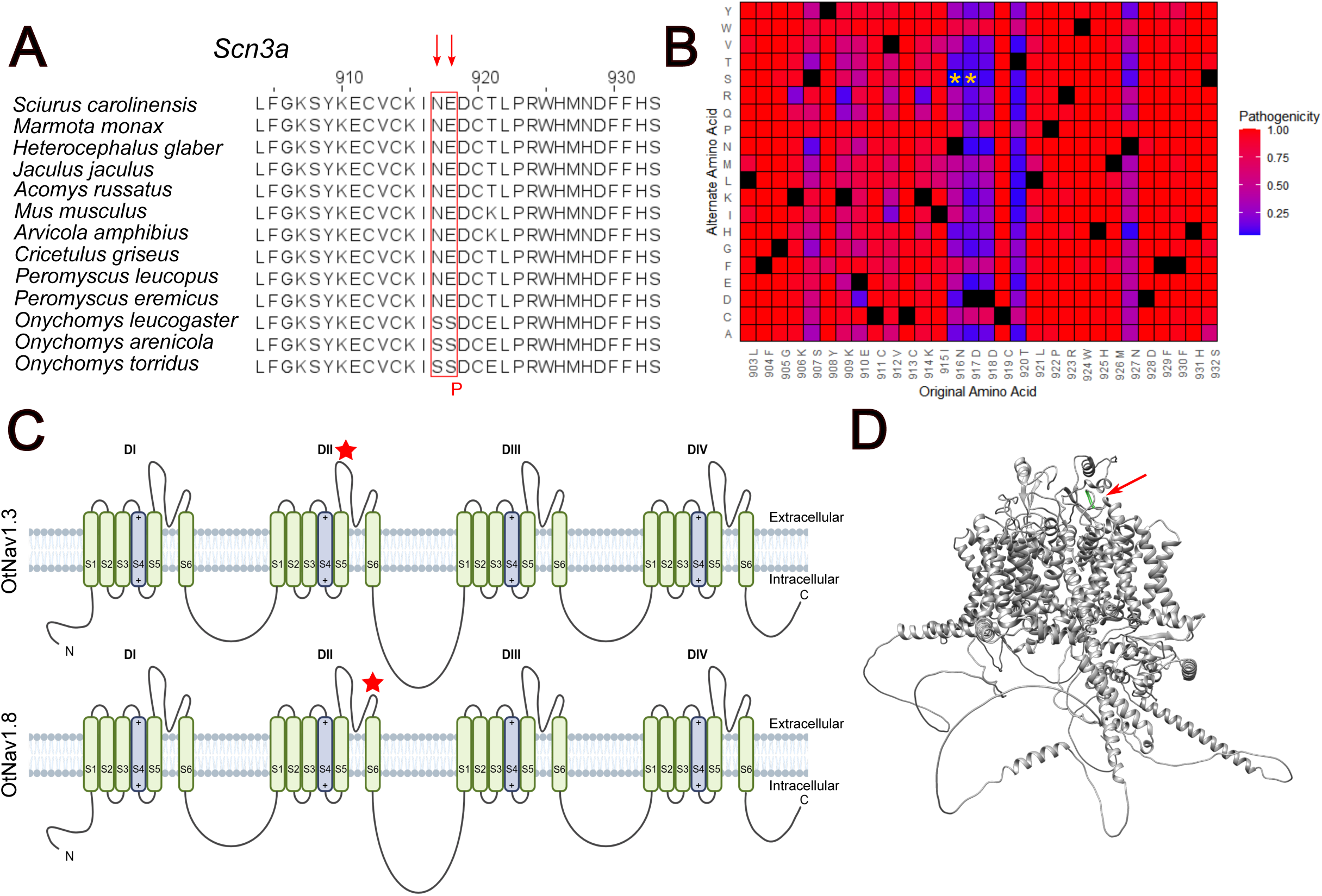
Positive selection in *Scn3a* in the *Onychomys* ancestral branch. **A)** Multiple sequence alignment of *Scn3a*. The positively selected amino acid sites are highlighted with a red box and arrow. A “P” below the second serine (S) indicates a predicted phosphorylation site. **B)** AlphaMissense heatmap (based on the human ortholog), showing predicted pathogenicity scores for all possible amino acid substitutions; asterisks denote substitutions found in *Onychomys*. **C)** Schematic diagrams of voltage-gated sodium channels: *Scn3a* (Nav1.3, **top**) and *Scn11a* (Nav1.8, **bottom**). A star marks the site under positive selection in Nav1.3, and a previously discovered amino acid substitution associated with toxin resistance is indicated in Nav1.8^16^. Created with BioRender.com **D)** AlphaFold-predicted structure of *Scn3a* in *O. torridus*; the positively selected amino acid is highlighted in green and marked with an arrow.

### Functional analysis

An organism’s response to toxin exposure involves a variety of gene pathways and protein interactions. Whilst it is known that sodium channel genes are critical for the transmission of pain signals from toxins, much less is currently known about the complete set of genes and proteins that act in concert to mediate the overall response to toxin exposure. A key question is whether the grasshopper mouse activates or deactivates proteins or their levels compared to susceptible species. We generated RNA-Seq data from sensory neuron tissue of mice exposed to toxic sprays from pinacate beetles. The tissue samples came from 30 mice from three species: *Onychomys torridus, Peromyscus eremicus,* and *Mus musculus. Peromyscus eremicus* coexist in the Sonoran Desert with grasshopper mice but do not prey on darkling beetles. *Mus musculus* are distantly related to grasshopper mice and do not prey on darkling beetles. Of the 30 mice, 15 were controls and had no exposure to beetle spray, and 15 mice were exposed to the beetle spray (Figure 5A). The mice were anaesthetised (not expected to affect sensory tissue gene expression) after exposure to the spray, and the tissue was extracted immediately. Mice were approximately two months old, and both male and female specimens were used for control and treatment groups. For each mouse, we generated RNA-Seq data from two different tissues: the dorsal root ganglion (DRG) and the trigeminal ganglion (TG). The DRG and TG are both key nerves for transmitting sensory signals such as touch, pain, and temperature to the central nervous system. The DRG are cell bodies of neurons that transmit signals from the body. The DRG are located just outside the spinal cord^34^. The TG is a cluster of nerve cell bodies for the trigeminal nerve (the fifth cranial nerve), responsible for sensation in the face ^35^.

**Figure 5:**
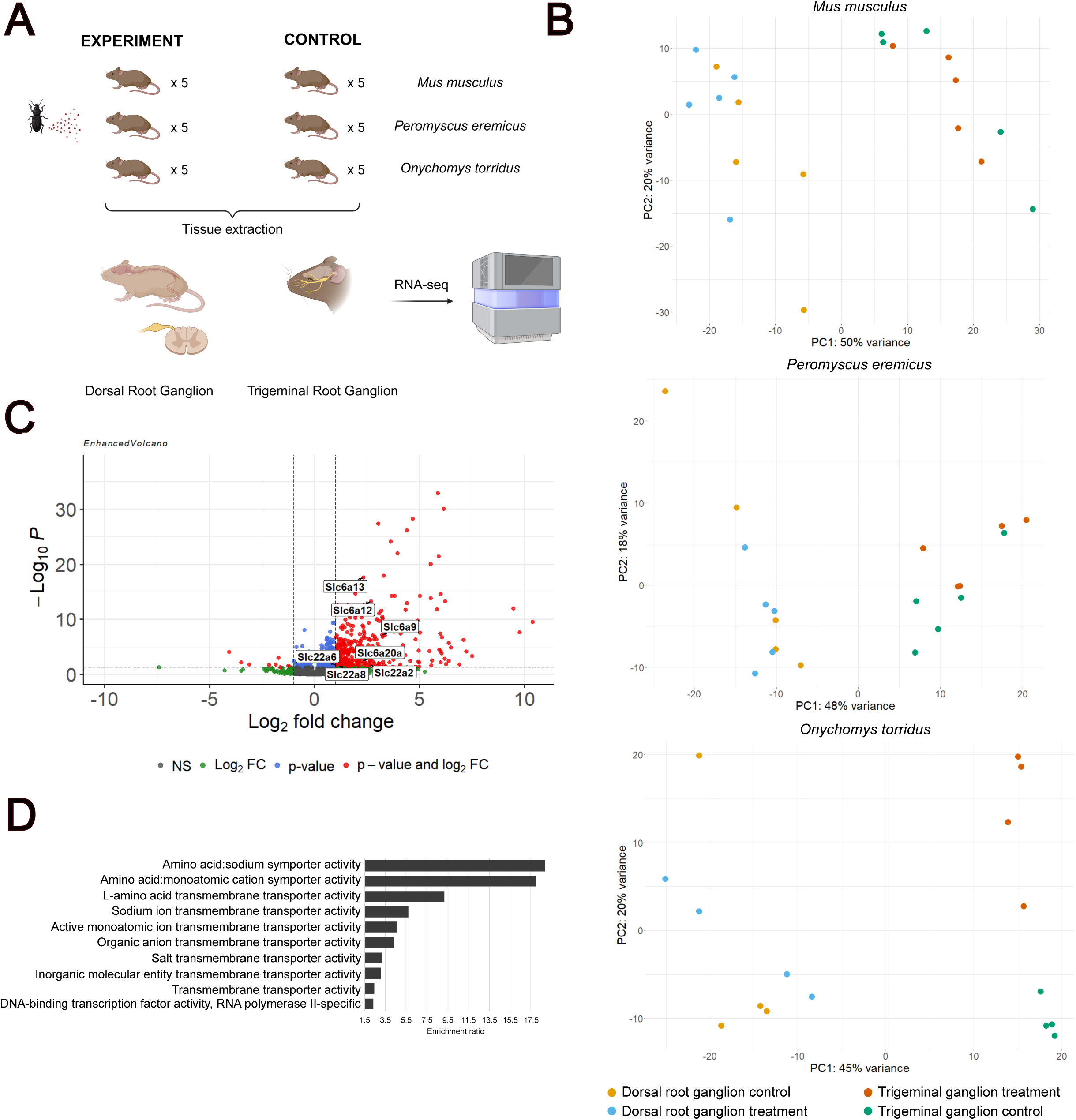
Pinacate beetle (*Eleodes*) toxin exposure experiment. **A)** Experimental design summary, including the number of species and tissues collected for RNA-seq before and after toxin exposure. Created with BioRender.com. **B)** Principal Component Analysis (PCA) of gene expression in *M. musculus, P. eremicus* and *O. torridus* samples: trigeminal ganglion and dorsal root ganglion. **C)** Volcano plot of differential gene expression in the *O. torridus* trigeminal ganglion following toxin exposure. Upregulated *Slc6a* and *Slc22a* gene family members are highlighted. **D)** PANTHER overrepresentation analysis showing enriched molecular function terms among upregulated genes.

Our first observation was that the gene expression values clustered by tissue and not by treatment status, so we decided to study the tissues separately (Figure 5B). We found that the data was clustered by treatment status in *O. torridus* in the trigeminal ganglion tissue, with no apparent clustering for the dorsal root ganglion. There was no treatment status clustering for any tissue in *P. eremicus* and *M. musculus* (Figure 5B). For *O. torridus* trigeminal ganglion samples, there were 354 genes differentially expressed with an adjusted p-value <0.05 (Benjamini–Hochberg test) and a Log_2_FC>1; 345 upregulated and 9 downregulated (Figure 5C and Supplementary Table 10). We did an enrichment test for all the upregulated genes, and 48.7% of the enriched genes are involved in transporter activity (Figure 5D). The *Slc22a* and *Slc6a* gene families (highlighted in Figure 5C) had several members upregulated.

## Discussion

Rodents have adapted and thrived in environmental niches, resulting in novel traits not found in laboratory mice that can be relevant in biomedical research. A notable example is the grasshopper mouse, which has developed toxin resistance to one of the most venomous scorpions in America, the AZ bark scorpion. The completion of high-quality reference genomes for the grasshopper mouse is a key component for establishing the *Onychomys* species as a model system for studying complex sensory phenotypes comprising toxin resistance and modified pain responses that, in turn, impact behavioural traits like predation and feeding. Here, we generated the first high-quality chromosome-scale genome assemblies, annotations and a pangenome for the *Onychomys* species and one of their closest species, *Peromyscus eremicus*.

We employed a pangenome approach to identify the most variable regions in the *Onychomys* genome. We found that most protein-coding genes in these regions were associated with defence and immunity functions, including members of the *Clec4* gene family. This pattern is consistent with previous observations in *M. musculu*s^36,37^. We detected an *Onychomys*-specific tandem gene duplication of the *Cblif* gene, which encodes a glycoprotein crucial for vitamin B12 absorption. Vitamin B12 comes from foods of animal origin, and it is essential for the health of nerve tissue, brain function, and red blood cells. Moreover, grasshopper mice have a modified stomach morphology due to their dietary specialisation, where parietal cells that express *Cblif* are especially numerous^7^. Interestingly, rodents from the genus *Muridae* have lost the *Tcn1* gene, which encodes for a transcobalamin^38^. Examination of several assemblies from *Criceitae* species revealed the same loss. *Tcn1* is present in other mammals and plays a role in the absorption of vitamin B12; it has been proposed to be an evolutionary product of a *Cblif* duplication^39^. In mammals, the *Tcn1* product, an haptocorrin (HC), binds to vitamin B12 in the stomach and is subsequently degraded by pancreatic enzymes, allowing *Cblif to* bind the vitamin and transport it to the ileum for absorption (Figure 2E). The grasshopper mouse does not have *Tcn1*; however, *Cblif* is duplicated, which may compensate for *Tcn1* loss, given its vitamin B12-rich carnivorous diet.

Our positive selection analysis identified that in *O. torridus,* residues Lys[107], Ile[126], and Arg[132] of *Foxe3* were positively selected. This gene encodes an ocular lens-specific transcription factor that plays an essential role in vertebrate lens formation and epithelial cell proliferation^31,32^. However, the mutations identified in this gene are heterozygous across the *O. torridus* population or homozygous for the ancestral allele. Comparative analysis of population data across the other *Onychomys* species revealed that while this gene is highly variable in *O. torridus*, it is in the ancestral state in *O. arenicola* and *O. leucogaster*. This observation may indicate an ongoing transition in the *O. torridus* population or suggest the presence of a heterozygous advantage, where individuals with heterozygous genotypes have a fitness advantage over those with either homozygous genotype. This may be due to *O. torridus* also preying on pinacate beetles, which release toxic sprays in the face and eyes of the grasshopper mouse during attacks^17^. Pinacate beetle spray contains benzoquinones, which target TRPA1 sensory transduction channels expressed in corneal tissue^18–21^. *Foxe3* plays an essential role during eye development, which likely explains why all individuals retain at least one ancestral copy. Nevertheless, the alternative alleles occur at a frequency of 17.5%; in a population of 20 individuals, 7 carried the alternative alleles in heterozygosity. These observations suggest that the alternative allele may contribute to the protection of the *O. torridus* eye lenses.

Rowe *et al.* found that Nav1.8 in *Onychomys torridus* has amino acid variants that bind scorpion venom peptides, which inhibit the Na+ currents and block action potential propagation to induce analgesia^16^. The characterised mutation consists of an inversion of the positions of a glutamine (Q859) and a glutamic acid (E862) residue in the domain II (DII) compared to corresponding residues in *M. musculus*. The position of the glutamic acid is essential for binding venom peptides that reduce Nav1.8 activity to block Na+ currents and pain signals in *O. torridus*. However, the Q861E replacement in *M. musculus* Nav1.8 did not completely block Na+ current, which indicates that although this mutation is necessary to inhibit Nav1.8 in *O. torridus*, it is likely not the only amino acid variant in DII contributing to venom sensitivity. We found that the mutation is also present in *Peromyscus eremicus.* While *P. eremicus* shares the same habitat as bark scorpions, they are not known to prey on the scorpions or to be resistant to the venom. These results suggest that *P. eremicus* has evolved some pain resistance mutations, and additional molecular adaptations may contribute to toxin pain resistance. Moreover, the E862 mutation has been found in other rodent and mammal species. This suggests that even though E862 may not have explicitly evolved due to selection by scorpion venom, *O. torridus* can use it to decrease its sensitivity to venom-induced pain.

We identified that residues Ser[867] and Ser[868] located in the extracellular part of the pore of domain II of Nav1.3 (sodium channel 3A) were under positive selection. This gene is expressed in the central nervous system (CNS) and is known to be involved in the generation and conduction of action potentials in excitable cells^40^. Additionally, it has been identified as a key player in nociceptive signalling, alongside Nav1.7, Nav1.8 and Nav1.9 ^40,41^, suggesting that mutations in this gene may contribute to pain resistance observed in *Onychomys* species. However, the gene is expressed in the CNS, which the venom cannot reach. Under normal conditions, peripheral nerve injury silences the mature pain system (Nav1.8 and Nav1.9) and reactivates a developmental hyperexcitable system (Nav1.3)^41,42^. The grasshopper mouse frequently engages in fights with other species; therefore, in the event of injury, the upregulation of Nav1.3 could confer resistance.

In our functional analysis, we anticipated that genes associated with cell repair might be upregulated, while those involved in sensory inflammation would be downregulated in *O. torridus*. However, what we observed was the upregulation of genes involved in transportation only in *O. torridus*, and notably, the *Slc22a* family had three genes

upregulated; this gene family encodes transporters that expel toxins from cells^43^. Moreover, the *Slc6a* family also had several members that were upregulated, encoding sodium-dependent neurotransmitter symporters ^44^. This gene family is expressed in the nervous system^45^. These findings suggest that *O. torridus* may expel toxins from its cells rapidly enough to confer resistance, a mechanism not observed in *P. eremicus* or *M. musculus*, which are susceptible. In susceptible species, we conclude that the transport response may be relatively weak or delayed, allowing intracellular toxin accumulation and subsequent cellular damage. By contrast, resistant species exhibit a rapid and robust transport response that facilitates efficient toxin export and therefore prevents cellular injury.

This study successfully assembled the first high-quality reference genomes and annotations for *Onychomys* species and *Peromyscus eremicus*, a key step in establishing *Onychomys* species as a model system for studying toxin and pain response, as well as behavioural traits. These genomes provide a foundational resource for future genomic research on these species. We detected an *Onychomys*-specific tandem gene duplication of the *Cblif* gene, an adaptation that likely supports the species’ dietary specialisation and modified stomach morphology. Additionally, we implemented a comprehensive pipeline to detect positive selection across genome-scale datasets. This led to the identification of a promising candidate gene for pain resistance in *Onychomys*: *Scn3a;* and *Foxe3* in *O. torridus,* which might be involved in lens protection against toxic sprays. Furthermore, we conducted functional analyses to gain a deeper understanding of gene regulation following toxin exposure. We observed the upregulation of genes associated with transportation in resistant species only, which may help expel toxins from the cell more quickly compared to susceptible species.

## Materials and Methods

### De novo reference genomes for *O. arenicola, O. leucogaster* & *P. eremicus*, and improved genome assembly for *O. torridus*

To assemble the genomes, we followed the Vertebrate Genomes Project (VGP) assembly pipeline. We assembled the PacBio HiFi reads into contigs using Hifiasm (Cheng et al., 2021), followed by quality control using assembly stats, BUSCO^46^ and Merqury^47^. Then we used purge_dups^48^ to remove haplotigs and false contig overlaps in the *de novo* assembly. Next, we used Hi-C data to scaffold the contigs into chromosome-scale super-scaffolds using Salsa2^49^ and YaHS^50^. We retained the Yahs scaffolds. The combination of HiFi reads with Hi-C resulted in near-complete partially phased chromosomes.

We carried out a round of manual curation following the ArimaPipeline (https://github.com/ArimaGenomics/CHiC) that aligns the Hi-C reads to the scaffolds and PretextMap, which converts these alignments into genomic contact maps that are read by PretextView, a desktop application for viewing and editing HiC contact maps. With the Hi-C contact maps, we were able to fix misassemblies, detect inversions, and anchor contigs to scaffolds. We then used a previously released version of the genome of *O. torridus* (GCA_903995425.1) to assign chromosomes in *O. arenicola* and *O. leucogaster* by aligning the genomes using MUMmer4^51^ and Gnuplot for visualisation. For *P. eremicus* we used the genome of *P. leucopus,* which is its closest species with a long-read-based available genome.

We ran FCS-GX, part of NCBI’s Foreign Contamination Screen (FCS) tool suite for the DNA contamination check^52^. To assess the abundance of different repeat elements, we ran RepeatMasker^53^ v4.1.4 with rmblastn v2.11.0+ and the Dfam 3.6 database, assuming the query species to be a rodent. Repeats were extracted from the .tbl output file generated by RepeatMasker.

### Genome Annotation

The gene annotations for the *Peromyscus* and *Onychomys* genome assemblies were produced using the Ensembl Gene Annotation pipeline^54^. The primary annotations were generated by aligning short-read transcriptomic datasets, sourced from the European Nucleotide Archive (ENA), to the respective genomes.

To address regions where transcript-based evidence was incomplete, additional annotation support was incorporated by aligning mammalian SwissProt proteins from UniProt^24^, which represent experimentally verified and curated protein sequences. Complementary evidence was also provided by mapping mouse coding sequences from the GENCODE resource^25^ through pairwise genome alignments.

Evidence from these sources was merged at each locus, with transcriptomic data given the highest priority when constructing the final gene models and the associated non-redundant transcript set. To distinguish functional isoforms from artefacts, open reading frames (ORFs) were compared against known vertebrate proteins, and models indicative of fragmentation or poor quality were filtered out. In genomic regions with sparse transcriptomic support, homology-based evidence was preferentially used, particularly when supported by strong intron-exon structure inferred from short-read data.

The resulting gene models were classified into three major categories: protein-coding genes, pseudogenes, and long non-coding RNAs (lncRNAs). Models with strong alignment to known proteins and without significant structural defects were designated as protein-coding. By contrast, models that matched proteins but displayed structural issues, such as missing initiation codons, abnormal splice sites, very short introns (<75 bp), or extensive repeat content, were reclassified as pseudogenes. Processed pseudogenes (retrotransposed copies) were identified as single-exon models with multi-exon homologs elsewhere in the genome.

Candidate lncRNAs were defined as models derived from transcriptomic data that did not overlap protein-coding regions and failed to meet the criteria for the other categories. However, single-exon transcripts were excluded from the lncRNA set, given their limited reliability.

Putative microRNAs (miRNAs) were predicted by aligning sequences from miRBase^55^ against the genomes, followed by RNAfold^56^ analysis of secondary structures. Other classes of small non-coding RNAs were identified using Rfam^57^, as described in^54^ PMID: 27337980, with structural validation performed through Infernal^58^.

The complete genome annotations for *Peromyscus* and *Onychomys* can be accessed through the updated Ensembl platform at beta.ensembl.org.

Ensembl’s gene predictions are based on human gene models, resulting in human gene names being assigned to the *Onychomys* and *Peromyscus* species. Therefore, we used Liftoff^59^ and Bedtools^60^ to map gene names from previous NCBI annotations to Ensembl annotations, converting them to the corresponding *Mus musculus* ortholog. Gene filtering was performed in five rounds, adjusting the Bedtools intersect parameter (-f), which specifies the minimum overlap required, from 0.9 to 0.1, decreasing by 0.2 in each round. After each round, genes that did not meet the current overlap threshold were retained and re-analysed with Bedtools intersect using the next, lower threshold. For each Ensembl gene, we retained the alias from the NCBI annotation.

### Synteny breaks detection for *O. torridus* and *P. eremicus*

*O. torridus* (GCA_949787125.1) and *P. eremicus* (GCA_949786415.1) genomes were aligned using Minimap2^61^ -asm10. Syri^62^ was used to detect synteny breaks, and Plotsr for visualisation^63^.

### Pangenome

Minigraph^26^ was used to produce the *Onychomys* pangenome. The *O. torridus* genome was used as the reference genome, and *O. arenicola* and *O. leucogaster* were aligned against it using the default parameters. This produced a graph composed of chains of bubbles, with the reference (*O. torridus*) as the backbone. Each bubble represents a structural variation. Gfatools (https://github.com/lh3/gfatools) was used to call the structural variations. From the structural variation file corresponding to each species, the chromosome and coordinates of the bubbles were extracted, along with the length of each bubble. These files, along with the length of *O. torridus* chromosomes (used as the backbone), were given to CIRCOS^64^ for visualisation. CIRCOS generated a histogram in which the X-axis represents chromosome positions and the Y-axis shows the amount of non-reference sequence across each chromosome in the *Onychomys* species. The Y-axis was scaled from 800bp to 200 Kbp.

### Most variable loci detection

To identify highly variable genomic regions in the Onychomys species pangenome, we extracted relevant coordinates and variation data from the species-level BED file generated by minigraph. From this file, only columns 1, 2, 3, and 8 were retained, corresponding to chromosome, start, end, and the length of the largest variation bubble (a measure of sequence divergence), respectively. A genome-wide sliding-window BED file (200 kb windows, 10 kb step) was generated spanning the O. torridus reference genome (used as the backbone in Minigraph). Using bedtools map, the windowed BED file and the filtered minigraph BED file were combined to generate a new BED file containing the amount of sequence divergence within each window. A custom Python script identified divergence-based loci by comparing consecutive windows; when the divergence difference exceeded a user-defined threshold (default: 50%), a new locus was defined. Locus coordinates were remapped to determine total sequence divergence, and the resulting file was sorted by divergence magnitude. The top 5% of windows showing the highest non-reference variation were retained, and bedtools intersect was used to identify and extract gene names that entirely overlapped (-f 1) these highly variable regions based on the O. torridus annotation file.

### Positive selection

Orthofinder^65^ was run using the proteomes of 14 mouse species (Supplementary Table 6), which were extracted from Ensembl. The selected species were those with chromosome-scale genome assemblies. We extracted one-to-one orthologs that were present in at least 13 species. The protein sequences of these genes were then aligned using ClustalO^66^. Using the corresponding transcript sequences and the protein alignments, codon alignments were generated with PAL2NAL^67^, filtering out any gaps. From the PAL2NAL alignments, we constructed phylogenetic trees using IQ-TREE2^68^. Finally, we ran PAML^29^ to detect positively selected genes, followed by a Benjamini and Hochberg correction for multiple testing. To further analyse the candidate genes, we developed a script that generated a table listing the gene names under positive selection, alignment lengths, and the amino acids positively selected based on a Bayes Empirical Bayes (BEB)^69^ analysis with probabilities greater than 0.95. From this table, we manually filtered out genes showing consecutive stretches of amino acids flagged as positively selected, as these were likely due to annotation or alignment errors. We also excluded genes with selected amino acids located at the very beginning or end of the alignment. This filtering process allowed us to focus on the most promising candidates, which were then manually reviewed. As a validation step, codeml was re-run for the candidate genes using the four-fold degenerate site species tree to check the robustness of the positive selection signal. This was not treated as an independent discovery stage, and no additional multiple-testing correction was applied.

### Illumina population data

The procedures used in the whole genome analyses were approved by the Institutional Animal Care and Use Committee (IACUC) at the following universities: The University of Texas at Austin AUP-2011-00103; Michigan State University AUP #10/13-229-00 and AUP #11/16-191-00; University of Oklahoma AUP #R21-015. Specimens of *O. torridus*, *O. arenicola*, *O. leucogaster*, and *P. eremicus* were trapped from locations in AZ, NM, and KS. Mice were euthanized using isoflurane before tissue extraction. Tissue samples were harvested from either wild caught mice or mice born to wild caught females. *O. torridus* were collected from two populations: the SRER (Santa Rita Experimental Range) AZ; and the Chiricahua Mountains, AZ. *O. arenicola* were collected from the Organ Mountains, NM. *O. leucogaster* were collected from a sand sage prairie habitat owned by the Sunflower Electric Power Corporation, KS. *P. eremicus* were collected from the SRER (Santa Rita Experimental Range) AZ. DNA samples were extracted from tissues using the Qiagen DNeasy DNA extraction kit. DNA samples were eluted in the AE buffer that came in the Qiagen DNeasy DNA extraction kit. Manufacturer’s guidelines for extraction were followed according to the Qiagen DNeasy DNA kit.

We generated Illumina population data for 22 *O. torridus*, 10 *O. arenicola*, 4 *O. leucogaster* and 11 *P. eremicus* individuals. The Illumina data was aligned against the reference genomes using BWA-mem2^70^ and then marked the duplicates using Samtools markdup. For quality control, we ran Samtools stats. All but three of our samples passed the QC check (Supplementary Table 8). To confirm no inter-species mislabeling of any of the samples, we compared the sequence for *Scn4a* (Sodium channel 1.4) from all our samples and made a phylogenetic tree (Figure 8) using IQ-TREE and FigTree that clustered the different species as expected. We ran Bcftools 1.16 for variant calling.

### RNA-Seq data

The experimental procedures used in the DGE analyses were approved by the Institutional Animal Care and Use Committee (IACUC) at the University of Oklahoma (Animal Use Protocol #R21-015). Sensory neurons (trigeminal ganglia and dorsal root ganglia) tissue were harvested (post euthanasia) from three species of mice (*Onychomys torridus*, *Peromyscus eremicus*, and *Mus musculus domesticus*). A total of ten individuals from each species were used for the experiments. Five mice were exposed to beetle spray and five control mice were exposed to distilled water. All procedures were approved by the University of Oklahoma Institutional Animal Care and Use committee (IACUC protocol #R21-015). Southern grasshopper mice (*Onychomys torridus*) and Cactus deer mice (*Peromyscus eremicus*) used in the DGE study were born in captivity (bred from wild caught females and males collected from the SRER, Santa Rita Experimental Range in southern AZ. House mice (*Mus musculus domesticus*, strain C57BL/6J) were ordered from The Jackson Laboratory, Bar Harbor, ME. All mice were anesthetized with a mixture of ketamine and xylazine. Beetle spray was collected from 50 individuals of darkling beetle (*Eleodes longicollus*). Aliquots of the spray mixture were placed in sterile atomizers. Distilled water was used as the control. Anesthetized mice were sprayed in the face with either beetle spray or distilled water. Mice were immediately euthanized. Trigeminal ganglia and dorsal root ganglia were harvested and flash frozen on dry ice. Samples were stored in a −80 degree C freezer until RNA extraction. Messenger RNA (mRNA) was extracted from tissue samples using Trizol protocol and stored in −80 degree C freezer.

RNA-Seq reads were mapped to the genomes using STAR^71^. For quality control, we ran Samtools stats (Supplementary Table 9). For the differential expression analysis, we used the STAR-featurecounts-Deseq2 pipeline^72,73^.

### Data Availability

The genome sequencing reads and the genome assemblies have been deposited at the European Nucleotide Archive under project accession: PRJEB60916. Table 1 contains the individual genome assembly accessions. The genomes and annotation are available from the Ensembl genome browser (https://beta.ensembl.org).

## References

1. Solari, S. & Baker, R. J. Mammal species of the world: a taxonomic and geographic reference. J. Mammal. 88, 824–830 (2007).

2. Fabre, P.-H., Hautier, L., Dimitrov, D. & Douzery, E. J. P. A glimpse on the pattern of rodent diversification: a phylogenetic approach. BMC Evol. Biol. 12, 88 (2012).

3. Seifert, A. W. et al. Skin shedding and tissue regeneration in African spiny mice (Acomys). Nature 489, 561–565 (2012).

4. Weinstein, S. B., Malanga, K. N., Agwanda, B., Maldonado, J. E. & Dearing, M. D. The secret social lives of African crested rats, Lophiomys imhausi. J. Mammal. 101, 1680–1691 (2020).

5. Rowe, A. H. & Rowe, M. P. Physiological resistance of grasshopper mice (Onychomys spp.) to Arizona bark scorpion (Centruroides exilicauda) venom. Toxicon 52, 597–605 (2008).

6. Campbell, P. et al. Vocal divergence is concordant with genomic evidence for strong reproductive isolation in grasshopper mice (Onychomys). Ecol. Evol. 9, 12886–12896 (2019).

7. Horner, B. E., Taylor, J. M. & Padykula, H. A. Food habits and gastric morphology of the grasshopper mouse. J. Mammal. 45, 513 (1964).

8. Rowe, A. H. & Rowe, M. P. Predatory grasshopper mice. Curr. Biol. 25, R1023–R1026 (2015).

9. Rodríguez de la Vega, R. C. & Possani, L. D. Overview of scorpion toxins specific for Na+ channels and related peptides: biodiversity, structure-function relationships and evolution. Toxicon 46, 831–844 (2005).

10. Rodríguez de la Vega, R. C. & Possani, L. D. Current views on scorpion toxins specific for K+-channels. Toxicon 43, 865–875 (2004).

11. Rowe, A. H. et al. Isolation and characterization of CvIV4: a pain inducing α-scorpion toxin. PLoS One 6, e23520 (2011).

12. Rowe, A. H. & Rowe, M. P. Risk assessment by grasshopper mice (Onychomys spp.) feeding on neurotoxic prey (Centruroides spp.). Anim. Behav. 71, 725–734 (2006).

13. Isbister, G. K. & Bawaskar, H. S. Scorpion envenomation. N. Engl. J. Med. 371, 457–463 (2014).

14. Stahnke, H. L. Some observations of the genus Centruroides marx (Buthidae, Scorpionida) and C. sculpturatus Ewing. Entomol. News 82, 281–307 (1971).

15. Parigi, A. A. et al. Complex physiological phenotypes: Structural variation in the muscle voltage-gated sodium channel imparts resistance to lethal scorpion toxins. SSRN Electron. J. (2021) doi:10.2139/ssrn.3940634.

16. Rowe, A. H., Xiao, Y., Rowe, M. P., Cummins, T. R. & Zakon, H. H. Voltage-gated sodium channel in grasshopper mice defends against bark scorpion toxin. Science 342, 441–446 (2013).

17. Eisner, T. Beetle’s spray discourages predators. Nat. Hist. 75, 42–47 (1966).

18. Macpherson, L. J. et al. Noxious compounds activate TRPA1 ion channels through covalent modification of cysteines. Nature 445, 541–545 (2007).

19. Ibarra, Y. & Blair, N. T. Benzoquinone reveals a cysteine-dependent desensitization mechanism of TRPA1. Mol. Pharmacol. 83, 1120–1132 (2013).

20. Talavera, K. et al. Mammalian transient receptor potential TRPA1 channels: From structure to disease. Physiol. Rev. 100, 725–803 (2020).

21. Okada, Y. et al. Transient receptor potential channels and corneal stromal inflammation. Cornea 34 **Suppl 11**, S136–41 (2015).

22. Kelly, T. S., Martin, R. A., Ronez, C., Cañón, C. & Pardiñas, U. F. J. Morphology and genetics of grasshopper mice revisited in a paleontological framework: reinstatement of Onychomyini (Rodentia, Cricetidae). J. Mammal. 104, 3–28 (2023).

23. Miller, J. R. & Engstrom, M. D. The Relationships of Major Lineages within Peromyscine Rodents: A Molecular Phylogenetic Hypothesis and Systematic Reappraisal. J Mammal 89, 1279–1295 (2008).

24. UniProt Consortium. UniProt: a worldwide hub of protein knowledge. Nucleic Acids Res. 47, D506–D515 (2019).

25. Frankish, A. et al. GENCODE reference annotation for the human and mouse genomes. Nucleic Acids Res. 47, D766–D773 (2019).

26. Li, H., Feng, X. & Chu, C. The design and construction of reference pangenome graphs with minigraph. Genome Biol. 21, 265 (2020).

27. Uto, T. et al. Clec4A4 is a regulatory receptor for dendritic cells that impairs inflammation and T-cell immunity. Nat. Commun. 7, 11273 (2016).

28. Alpers, D. H. & Russell-Jones, G. Gastric intrinsic factor: the gastric and small intestinal stages of cobalamin absorption. a personal journey. Biochimie 95, 989–994 (2013).

29. Yang, Z. PAML 4: phylogenetic analysis by maximum likelihood. Mol. Biol. Evol. 24, 1586–1591 (2007).

30. Blixt, A. et al. A forkhead gene, FoxE3, is essential for lens epithelial proliferation and closure of the lens vesicle. Genes Dev. 14, 245–254 (2000).

31. Brownell, I., Dirksen, M. & Jamrich, M. Forkhead Foxe3 maps to the dysgenetic lens locus and is critical in lens development and differentiation. Genesis 27, 81–93 (2000).

32. Semina, E. V., Brownell, I., Mintz-Hittner, H. A., Murray, J. C. & Jamrich, M. Mutations in the human forkhead transcription factor FOXE3 associated with anterior segment ocular dysgenesis and cataracts. Hum. Mol. Genet. 10, 231–236 (2001).

33. Cheng, J. et al. Accurate proteome-wide missense variant effect prediction with AlphaMissense. Science 381, eadg7492 (2023).

34. Berger, A. A. et al. Dorsal root ganglion (DRG) and chronic pain. Anesth. Pain Med. 11, e113020 (2021).

35. Messlinger, K. & Russo, A. F. Current understanding of trigeminal ganglion structure and function in headache. Cephalalgia 39, 1661–1674 (2019).

36. Lilue, J. et al. Sixteen diverse laboratory mouse reference genomes define strain-specific haplotypes and novel functional loci. Nat. Genet. 50, 1574–1583 (2018).

37. Lilue, J., Shivalikanjli, A., Adams, D. J. & Keane, T. M. Mouse protein coding diversity: What’s left to discover? PLoS Genet. 15, e1008446 (2019).

38. Hygum, K. et al. Mouse transcobalamin has features resembling both human transcobalamin and haptocorrin. PLoS One 6, e20638 (2011).

39. Mucha, P., Kus, F., Cysewski, D., Smolenski, R. T. & Tomczyk, M. Vitamin B12 metabolism: A network of multi-protein mediated processes. Int. J. Mol. Sci. 25, 8021 (2024).

40. de Lera Ruiz, M. & Kraus, R. L. Voltage-gated sodium channels: Structure, function, pharmacology, and clinical indications. J. Med. Chem. 58, 7093–7118 (2015).

41. Hains, B. C. et al. Upregulation of sodium channel Nav1.3 and functional involvement in neuronal hyperexcitability associated with central neuropathic pain after spinal cord injury. J. Neurosci. 23, 8881–8892 (2003).

42. Xu, W., Zhang, J., Wang, Y., Wang, L. & Wang, X. Changes in the expression of voltage-gated sodium channels Nav1.3, Nav1.7, Nav1.8, and Nav1.9 in rat trigeminal ganglia following chronic constriction injury. Neuroreport 27, 929–934 (2016).

43. Nigam, S. K. The SLC22 transporter family: A paradigm for the impact of drug transporters on metabolic pathways, signaling, and disease. Annu. Rev. Pharmacol. Toxicol. 58, 663–687 (2018).

44. Pramod, A. B., Foster, J., Carvelli, L. & Henry, L. K. SLC6 transporters: structure, function, regulation, disease association and therapeutics. Mol. Aspects Med. 34, 197–219 (2013).

45. Bröer, S. & Gether, U. The solute carrier 6 family of transporters: The Solute Carrier Family 6. Br. J. Pharmacol. 167, 256–278 (2012).

46. Tegenfeldt, F. et al. OrthoDB and BUSCO update: annotation of orthologs with wider sampling of genomes. Nucleic Acids Res. 53, D516–D522 (2025).

47. Rhie, A., Walenz, B. P., Koren, S. & Phillippy, A. M. Merqury: reference-free quality, completeness, and phasing assessment for genome assemblies. Genome Biol. 21, 245 (2020).

48. Guan, D. et al. Identifying and removing haplotypic duplication in primary genome assemblies. Bioinformatics 36, 2896–2898 (2020).

49. Ghurye, J., Pop, M., Koren, S., Bickhart, D. & Chin, C.-S. Scaffolding of long read assemblies using long range contact information. BMC Genomics 18, 527 (2017).

50. Zhou, C., McCarthy, S. A. & Durbin, R. YaHS: yet another Hi-C scaffolding tool. Bioinformatics 39, btac808 (2023).

51. Marçais, G. et al. MUMmer4: A fast and versatile genome alignment system. PLoS Comput. Biol. 14, e1005944 (2018).

52. Astashyn, A. et al. Rapid and sensitive detection of genome contamination at scale with FCS-GX. Genome Biol. 25, 60 (2024).

53. Chen, N. Using RepeatMasker to identify repetitive elements in genomic sequences. Curr. Protoc. Bioinformatics **Chapter** 4, Unit 4.10 (2004).

54. Aken, B. L. et al. The Ensembl gene annotation system. Database (Oxford) 2016, (2016).

55. Kozomara, A., Birgaoanu, M. & Griffiths-Jones, S. miRBase: from microRNA sequences to function. Nucleic Acids Res. 47, D155–D162 (2019).

56. Denman, R. B. Using RNAFOLD to predict the activity of small catalytic RNAs. Biotechniques 15, 1090–1095 (1993).

57. Kalvari, I. et al. Rfam 13.0: shifting to a genome-centric resource for non-coding RNA families. Nucleic Acids Res. 46, D335–D342 (2018).

58. Nawrocki, E. P. & Eddy, S. R. Infernal 1.1: 100-fold faster RNA homology searches. Bioinformatics 29, 2933–2935 (2013).

59. Shumate, A. & Salzberg, S. L. Liftoff: accurate mapping of gene annotations. Bioinformatics 37, 1639–1643 (2021).

60. Quinlan, A. R. & Hall, I. M. BEDTools: a flexible suite of utilities for comparing genomic features. Bioinformatics 26, 841–842 (2010).

61. Li, H. Minimap2: pairwise alignment for nucleotide sequences. Bioinformatics 34, 3094–3100 (2018).

62. Goel, M., Sun, H., Jiao, W.-B. & Schneeberger, K. SyRI: finding genomic rearrangements and local sequence differences from whole-genome assemblies. Genome Biol. 20, 277 (2019).

63. Goel, M. & Schneeberger, K. Plotsr: Visualizing structural similarities and rearrangements between multiple genomes. Bioinformatics 38, 2922–2926 (2022).

64. Krzywinski, M. et al. Circos: an information aesthetic for comparative genomics. Genome Res. 19, 1639–1645 (2009).

65. Emms, D. M. & Kelly, S. OrthoFinder: phylogenetic orthology inference for comparative genomics. Genome Biol. 20, 238 (2019).

66. Sievers, F. & Higgins, D. G. Clustal Omega, accurate alignment of very large numbers of sequences. Methods Mol. Biol. 1079, 105–116 (2014).

67. Suyama, M., Torrents, D. & Bork, P. PAL2NAL: robust conversion of protein sequence alignments into the corresponding codon alignments. Nucleic Acids Res. 34, W609–12 (2006).

68. Minh, B. Q. et al. IQ-TREE 2: New models and efficient methods for phylogenetic inference in the genomic era. Mol. Biol. Evol. 37, 1530–1534 (2020).

69. Yang, Z., Wong, W. S. W. & Nielsen, R. Bayes empirical bayes inference of amino acid sites under positive selection. Mol. Biol. Evol. 22, 1107–1118 (2005).

70. Li, H. & Durbin, R. Fast and accurate short read alignment with Burrows-Wheeler transform. Bioinformatics 25, 1754–1760 (2009).

71. Dobin, A. et al. STAR: ultrafast universal RNA-seq aligner. Bioinformatics 29, 15–21 (2013).

72. Liao, Y., Smyth, G. K. & Shi, W. featureCounts: an efficient general purpose program for assigning sequence reads to genomic features. Bioinformatics 30, 923–930 (2014).

73. Love, M. I., Huber, W. & Anders, S. Moderated estimation of fold change and dispersion for RNA-seq data with DESeq2. Genome Biol. 15, 550 (2014).

