## Supplementary material for "The *Onychomys* pangenome reveals the unique molecular adaptations that confer toxin resistance": Table 1

| Species | Assembly accession | Sex | Length (Gbp) | N bases (Mbp) | Protein coding genes | BUSCO (%) | SINE (Mbp) | LINE (Mbp) | ERVs (Mbp) |
|---|---|---|---|---|---|---|---|---|---|
| *Peromyscus eremicus H1* | GCA_949786435.1 | F | 2.41 | 5.304 | 20,373 | 92.8 | 221.64 | 290.28 | 362.86 |
| *Peromyscus eremicus H2* | GCA_949786415.1 | F | 2.38 | 5.09 | 20,223 | 93.1 | 222.03 | 291.26 | 363.00 |
| *Onychomys torridus* | GCA_949787125.1 | F | 2.4 | 244.31 | 19,182 | 93.8 | 215.39 | 310.94 | 355.54 |
| *Onychomys arenicola H1* | GCA_949786405.1 | F | 2.56 | 0.36 | 20,095 | 94.8 | 221.08 | 330.76 | 379.25 |
| *Onychomys arenicola H2* | GCA_949786425.1 | F | 2.51 | 0.32 | 19,091 | 93.5 | 215.71 | 321.20 | 367.18 |
| *Onychomys leucogaster H1* | GCA_949786395.1 | M | 2.37 | 0.282 | 19,018 | 92.9 | 213.37 | 318.08 | 357.97 |
| *Onychomys leucogaster H2* | GCA_949786385.1 | M | 2.5 | 0.35 | 19,661 | 94.2 | 219.32 | 334.35 | 375.62 |
