## Supplementary material for "The *Onychomys* pangenome reveals the unique molecular adaptations that confer toxin resistance": Table 2

| Branch | Gene name | Chromosome | Position | AA change |
|---|---|---|---|---|
| *Onychomys* ancestral | ***Scn3a*** | OncTor#Chr4 | 42,305,187-42,396,120 | N867S, E868S |
| *Onychomys* ancestral | *Hnrnph1*** | OncTor#Chr8 | 38,718,998-38,727,713 | S435E |
| *Onychomys* ancestral | *Itih3* | OncTor#Chr9 | 27,212,395-27,227,730 | P815G |
| *Onychomys* ancestral | *Rims2* | OncTor#Chr16 | 6,607,900-7,071,673 | Y320R, R324E |
| *Onychomys* ancestral | *Gpalpp1* | OncTor#Chr9 | 64,359,698-64,384,817 | S42R, D45S |
| *Onychomys* ancestral | *Irs1* | OncTor#Chr23 | 65,960,835-66,018,849 | P982S |
| *Onychomys* ancestral | *Dennd1a* | OncTor#Chr4 | 1,037,986-1,517,768 | L467K, K469V, E473S |
| *Onychomys* ancestral | *Spag5* | OncTor#Chr8 | 66,603,970-66,623,698 | N1000R |
| *Onychomys* ancestral | *Otud1** | OncTor#Chr5 | 78,676,808-78,679,871 | E243G |
| *Onychomys* ancestral | *Npas2* | OncTor#Chr18 | 34,083,215-34,262,263 | R570Q |
| *Onychomys* ancestral | ENSOTOG00010017238 | OncTor#Chr19 | 60,621,399-60,635,823 | A91I |
| *Onychomys torridus* | ***Foxe3*** | OncTor#Chr2 | 129,697,432-129,698,265 | R107K, V126I, N132R |
| *Onychomys torridus* | *Krt86* | OncTor#Chr16 | 66,301,762-66,307,671 | R24K |
| *Onychomys torridus* | *Akt1* | OncTor#Chr14 | 80,056,698-80,078,780 | K356R |
| *Onychomys torridus* | *Nanos1* | OncTor#Chr1 | 180,379,151-180,379,972 | E131S |
| *Onychomys torridus* | *Cabp2* | OncTor#Chr1 | 123,865,018-123,869,704 | R93T |
| *Onychomys torridus* | *Antxr1* | OncTor#Chr3 | 112,318,176-112,509,938 | Q136A |
| *Onychomys torridus* | *Bace1* | OncTor#Chr7 | 39,286,868-39,311,680 | S317G |
| *Onychomys torridus* | *Ubr3* | OncTor#Chr4 | 46,751,609-46,901,022 | K1798L |
| *Onychomys torridus* | *Sipa1l2* | OncTor#Chr5 | 90,566,208-90,651,831 | S437G |
| *Onychomys torridus* | *Acot11* | OncTor#Chr2 | 123,415,000-123,474,024 | K364R |
| *Onychomys torridus* | *Sfrp5* | OncTor#Chr1 | 160,707,335-160,712,329 | A94N, R96N, S99M |
| *Onychomys torridus* | *Cldn4* | OncTor#Chr22 | 18,363,105-18,364,940 | N53S |
| *Onychomys torridus* | *Dph2* | OncTor#Chr2 | 132,498,799-132,504,184 | L164A, A165C, A166L |
| *Onychomys arenicola* | *Zfyve1* | OncAre#1#Chr14 | 53,396,611-53,449,782 | K64G |
| *Onychomys arenicola* | *Retreg1* | OncAre#1#Chr15 | 63,894,567-63,927,820 | S174N |
| *Onychomys arenicola* | *Ctnnd1* | OncAre#1#Chr4 | 61,980,572-62,033,841 | A787C |
| *Onychomys arenicola* | *Asb3* | OncAre#1#Chr10 | 3,643,404-3,862,890 | S488N |
| *Onychomys arenicola* | *Slc16a2* | OncAre#1#ChrX | 81,197,606-81,319,452 | S291N |

| | | | | |
|---|---|---|---|---|
| *Onychomys leucogaster* | *Apoa1* | OncLeu#2#Chr7 | 39,210,075-39,211,780 | F246Q, R247D, M249H |
| *Onychomys leucogaster* | *A530016L24Rik* | OncLeu#2#Chr14 | 80,223,307-80,232,106 | P94D |
